# Early auditory and adult mating experiences interact with singer identity to shape neural responses to song in female zebra finches

**DOI:** 10.1101/2024.10.04.616742

**Authors:** Isabella Catalano, Sarah C. Woolley

## Abstract

Social and sensory experiences across the lifespan can shape social interactions, however, experience-dependent plasticity is widely studied within discrete life stages. In the socially monogamous zebra finch, in which females use learned vocal signals to identify individuals and form long-lasting pair bonds, developmental exposure to song is key for females to show species-typical song perception and preferences. While adult mating experience can still lead to pair-bonding and song preference learning even in birds with limited previous song exposure (“song-naïve”), whether similarities in adult behavioral plasticity between normally-reared and song-naïve females reflect convergent patterns of neural activity is unknown. We investigated this using expression of a marker of neural activity and plasticity (phosphorylated S6) in mated normally-reared and song-naïve females in response to song from either their mate, a neighbor, or an unfamiliar male. We found that, in portions of a secondary auditory region (the caudomedial nidopallium NCM) and in dopaminergic neurons of the caudal ventral tegmental area, hearing the mate’s song significantly increased pS6 expression in females from both rearing conditions. In contrast, within other NCM subregions, song-identity drove different patterns of pS6 expression depending on the rearing condition. These data suggest that developmental experiences can have long-lasting impacts on the neural signatures of behaviors acquired in adulthood and that socially-driven behavioral plasticity in adults may arise through both shared and divergent neural circuits depending on an individual’s developmental experiences.

## INTRODUCTION

Early sensory experiences adaptively shape perception as well as the structure and function of neural systems. Developmental sensory experience can have especially long-lasting effects on auditory perception and discrimination. For example, language exposure in infants leads to phonemic tuning whereby infants lose the ability to discriminate between a broad variety of sounds and become tuned to their own language(s) (see Moore & Linthicum, 2007). In oscine songbirds, juvenile exposure to song is similarly important. Classic studies in white-crowned sparrows found that juvenile birds will learn the ‘dialect’ of song they are exposed to, rather than the one of their genetic parents (Marler & Tamura, 1964). Early exposure to communication signals also tunes neural activity. For example, male zebra finches, who learn song from a tutor, have cells in a secondary auditory cortical region, the caudomedial nidopallium (NCM), that become selective for their tutor’s song over the course of development while males raised without tutor exposure do not show similar tuning (Yanagihara & Yazaki-Sugiyama, 2016), and produce species-atypical song in adulthood (Eales, 1985; Jones et al., 1996).

Song perception is also shaped by early exposure to song in female zebra finches. Even though female zebra finches do not learn to sing, juvenile females still form a lasting memory of the tutor’s song (Clayton, 1988; Riebel, 2000) and become tuned to the songs or dialect from their own colony (Le Maguer et al., 2021). Additionally, hearing song during development is critical for both auditory discrimination and species-typical song preferences in females. For example, compared to normally-reared females, female zebra finches who are raised without exposure to song (“song-naïve”) show impaired frequency discrimination abilities (Sturdy et al., 2001) and do not show species-typical preferences for courtship song over non-courtship song (Chen et al., 2017). Further, developmental experience also impacts the evoked neural responses to preferred and less-preferred songs. Unlike normally-reared females, song-naïve females do not show greater responses to courtship compared to non-courtship songs in either a secondary auditory pallial region (caudomedial nidopallium [NCM], Chen et al., 2017) or in dopaminergic neurons in the caudal ventral tegmental area (cVTA; Barr & Woolley, 2018). Thus, developmental sensory experience can dramatically impact sensory perception and neural responses throughout life.

While experience with conspecific song early in development shapes discrimination and species-typical preferences, adult social experiences can also lead to lasting changes to perception and preference in both song-naïve and normally-reared birds. Among adult songbirds, song is important to determine the species and identity of the singer and, in many species, is used in mate selection (see Nowicki & Searcy, 2005 for review). In zebra finches, males and females form lifelong pair-bonds (Riebel, 2009) and pair-bonded females strongly prefer to hear the song of their partner over the song of an unfamiliar bird (Miller, 1979; Wall & Woolley, 2024; Woolley & Doupe, 2008). Intriguingly, adult pair-bonding experience can drive song preference learning in song-naïve females and even promote species-typical preferences for unfamiliar courtship songs like those seen in normally-reared birds (Wall & Woolley, 2024). However, while song-naïve females form pair-bonds with a partner and preferences for his song, there were important differences. For example, the relationships between pair bonding behaviors and the strength of preference differed between normally-reared and song-naïve females, indicating that there could be differences in the way that birds learn to associate the song with particular social interactions. These data hint that while early exposure to song is not necessary for adult social experiences to drive preference learning, differences in developmental experience could lead to differences in the manner in which these preferences are formed and maintained, perhaps through differential effects on the neural substrates of song preferences.

Here, we investigated whether the ability of song-naïve birds to learn preferences for a partner’s songs was associated with similar patterns of activity throughout the brain, especially in the auditory forebrain and catecholamine producing neurons in the mid- and hindbrain, compared to normally-reared birds. We quantified expression of an activity-dependent neural marker, phosphorylated S6 (pS6), in pair-bonded females from both rearing conditions in response to songs with differing degrees of familiarity. Specifically, within a cohort, all females heard the same song, but each female had a different relationship to the singer: he was either their mate, a familiar neighbor, or an unfamiliar male. We found that the impact of rearing condition on the response to stimulus identity varied across regions, highlighting differential effects of mating experience on evoked activity depending on an individual’s developmental experience.

## RESULTS

For female zebra finches, developmental exposure to song is key for tuning auditory responses and the display of species-typical song preferences. Here, we investigated whether neural circuits important for learned song preferences in adult females are affected by exposure to song when birds are juveniles. We created experimental cohorts of normally-reared and song-naïve females (see Methods for details on rearing). For each cohort in each rearing condition, we housed a male-female pair in a cage within a sound-attenuating chamber (“soundbox”). Within the same soundbox, a second male-female pair was housed in a different, adjacent cage. Cages were separated by a visual barrier which allowed birds in each cage to hear but not see or physically interact with birds in the neighboring cage. We repeated this configuration across two soundboxes for each rearing condition. After two weeks of cohabitation, females were housed in isolation and all females within a cohort heard playback of the song of the same male or silence. Importantly, while all females heard the same song, because of the experimental design, each bird had a different relationship with the singer: he was either their partner, an acoustically familiar male (same soundbox), or an unfamiliar male (different soundbox; Fig. 1). We then quantified expression of pS6 catecholemine-producing cells in the mid- and hindbrain (caudal and rostral ventral tegmental area [cVTA, rVTA], substantia nigra pars compacta [SNc], periaqueductal grey [PAG], locus coeruleus [LC]; Fig1e, red in figures), in ascending (lateral mesencephalic nucleus [MLd], nucleus ovoidalis [Ov]; Fig. 1f, teal), primary (caudolateral mesopallium [CLM], Fig. 1f, green), and secondary auditory regions (caudomedial mesopallium [CMM], Fig. 1f, green; caudomedial nidopallium [NCM], Fig. 1f, blue), and two regions in the social behavior network, the lateral septum (LS) and ventromedial hypothalamus (VMH; Fig. 1f, orange). We found that the dopamine-producing cVTA as well as the auditory regions MLd, CMM, and NCM all showed significant increases in pS6 in birds that heard song compared to silence (cVTA: *X^2^* _(1, 39)_ = 5.3865, *p* = 0.0203; MLd: *X^2^* _(1, 39)_ = 9.9837, *p* = 0.0016; CMM: *X^2^* _(1, 39)_ = 21.4878, *p* <0.0001; NCM: *X^2^* _(1, 39)_ = 47.5448, *p* < 0.0001; all other regions *p* > 0.13; Fig. 1e-f). We focused on these song-responsive regions for further analysis.

**Fig. 1.**
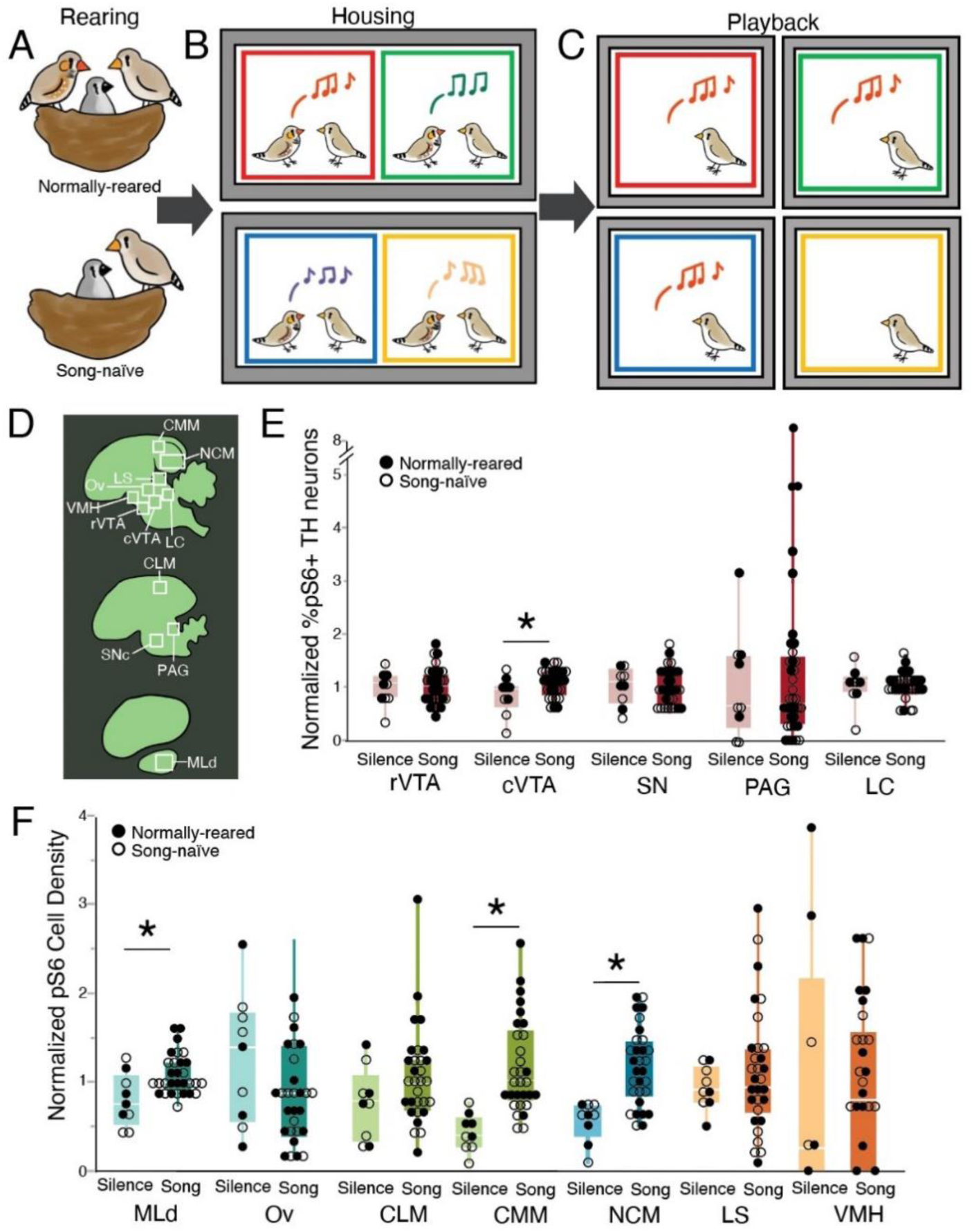
Song-evoked pS6 expression. (A) Females were reared either with both parents (normally-reared) or with just their mothers (song-naïve). (B) Then, as adults, females from both rearing conditions were housed for two weeks with a male partner. Two pairs of birds were housed within the same soundbox. (C) After two weeks, all birds were played the song of the same male, but birds differed in their relationship to that male: he was either their mate (red), a familiar neighbor (green) or unfamiliar (blue). A fourth group heard no songs (yellow). (D) Illustrations of sagittal sections of the avian brain (green) indicating the locations of pS6+ cell counts (white boxes) in catecholaminergic, auditory, and social behavior network regions. (E) Box and whisker plots of the normalized percent of tyrosine hydroxylase cells that co-localize with pS6 in catecholamine-positive regions and the (F) normalized density of pS6 expressing cells in auditory and SBN regions. Points are mean counts per individual for normally-reared (closed circles) and song-naïve females (open circles). Each box spans the interquartile range, horizontal white lines indicate the median and whiskers show the minima and maxima. * indicates a difference at p < 0.05.

### Mate’s song increases activation of dopaminergic neurons in females from both rearing conditions

Within the VTA, we quantified the number of active catecholaminergic neurons by counting both the number of pS6-immunoreactive cells (IR) and tyrosine hydroxylase immunoreactive cells (TH-IR), then calculating the percentage of active TH-IR cells (colocalization of TH-IR and pS6-IR). Based on previous work (Bambico et al., 2019; Barr & Woolley, 2018; Goodson et al., 2009; Lammel et al., 2008; Matheson & Sakata, 2015), we separated the VTA into rostral (rVTA) and caudal (cVTA) regions. In the cVTA there was a significant main effect of stimulus, (*X^2^* = 10.4291, *p* = 0.0152; Fig 2). Specifically, pS6 expression in dopaminergic neurons in the cVTA was significantly greater in birds that heard their mate’s song compared to those that heard an unfamiliar song (*p* = 0.0243) or silence (*p* = 0.0021). Females that heard familiar song had pS6 expression that was intermediate between those that heard mate’s song and those that heard unfamiliar song or silence (*p* > 0.09 for all comparisons). This effect was consistent across rearing conditions and parallels the similarity in song preference previously reported between normally-reared and song-naïve females (Wall & Woolley, 2024).

**Fig. 2.**
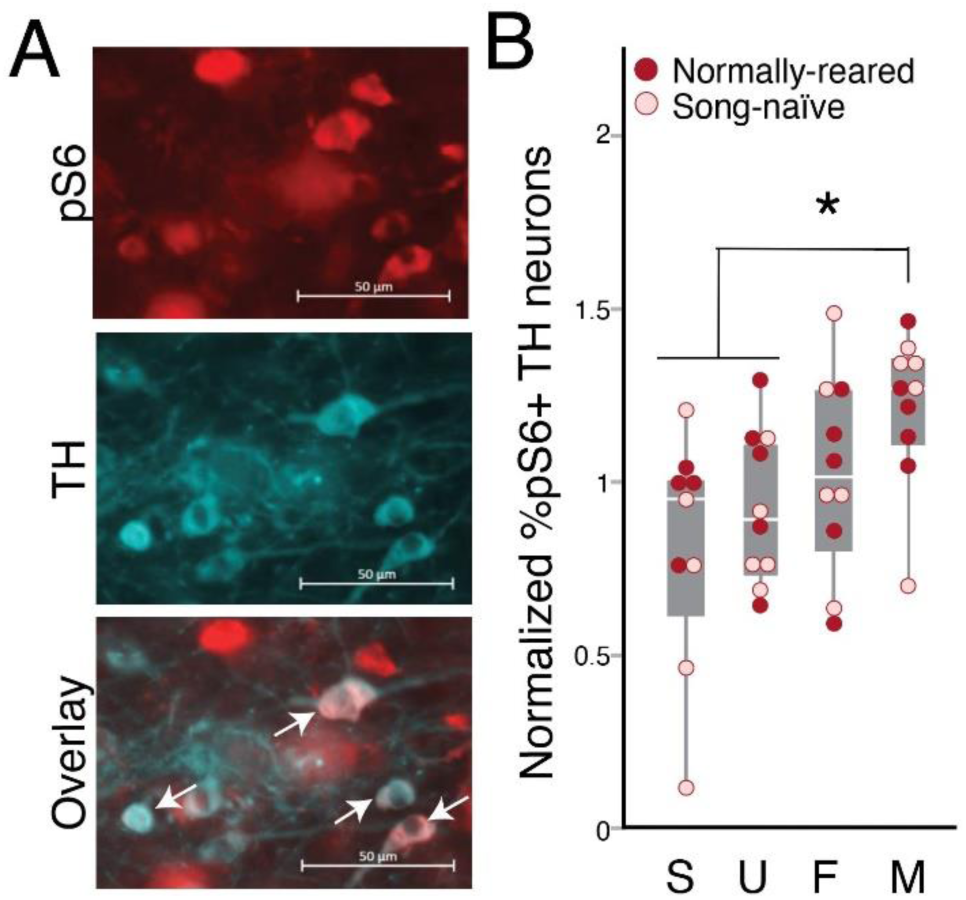
cVTA results.(A) Images of staining for pS6 in red (top), tyrosine hydroxylase in cyan (middle) and the overlay (bottom) in the cVTA. Arrows highlight cells expressing both pS6 and TH. (B) The percent of TH neurons expressing pS6 was significantly greater in females that heard their mate’s song compared to females that heard an unfamiliar song or silence. Points are normalized mean counts per individual for normally-reared (closed circles) and song-naïve females (open circles). * indicates a difference at p < 0.05.

### Mate’s song drives high levels of pS6 expression across the NCM in normally-reared females

The NCM is a broad region that encompasses much of the pallium caudal to the primary auditory region Field L. Previous work has found somewhat uniform induction of immediate early genes, such as EGR1, throughout the NCM. In contrast, pS6 expression in the NCM has been described as more irregular, with clusters of pS6+ cells within particular subregions (Ahmadiantehrani et al., 2018). We found similarly clustered expression of pS6 as has been previously described. Notably, we saw dense pS6 expression in a dorsocaudal region, an ventroanterior region, and in a central diagonal band that runs parallel to the lamina mesopallialis separating the meso and nidopallium and, more laterally, with the border of Field L3 (see Fig. 3a). We quantified pS6 expression in four locations within these regions that could be consistently identified using local landmarks in DAPI: one in the dorsocaudal population (NCMd), one in the ventroanterior region (NCMv), one centrally within the diagonal band of pS6 expression (NCMc), and one at the junction of the hippocampus and posterior NCM (NCMp; see Fig. 3b). Across all regions, the greatest increase in pS6 expression was generally in response to the mate’s song, while the lowest expression was in silent controls. However, there was also variation between regions in the level of pS6 induction in the response to stimulus identity (*X^2^* _39)_ = 22.2655, *p* = 0.0081) as well as significant interactions between stimulus identity and developmental song exposure (*X^2^* = 21.2266, *p* <0.0001).

**Fig. 3.**
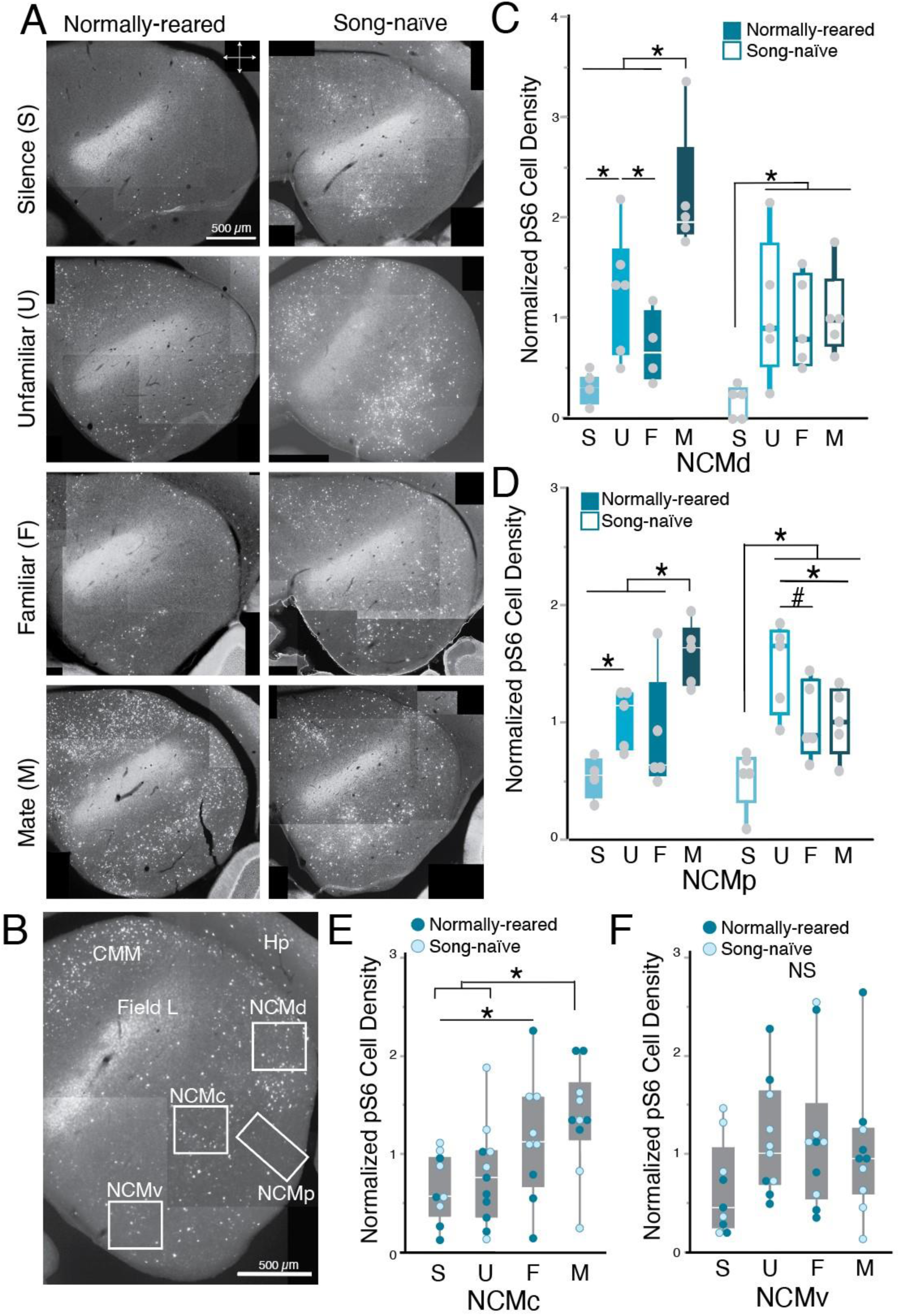
NCM results differ by subregion. (A) Representative photomicrographs of pS6 expression (white) throughout the caudal meso-and nidopallium for all 8 groups. (B) Photomicrograph of the caudal auditory forebrain. White boxes indicate the four regions that were counted within the NCM. (C-E) Box and whisker plots of pS6 expression varied between stimuli and rearing conditions in the (C) dorsal (NCMd), (D) posterior (NCMp), and (E) caudal (NCMc) NCM. (F) There were no significant effects of the stimulus or rearing condition on pS6 expression in the ventral NCM (NCMv). CMM Caudomedial mesopallium; Hp hippocampus; NCMc central caudomedial nidopallium; NCMd dorsal caudomedial nidopallium; NCMp posterior caudomedial nidopallium; NCMv ventral caudomedial nidopallium. Points are mean counts per individual for normally-reared (filled bars in C-D, closed circles in E-F) and song-naïve females (open bars in C-D, open circles in E-F). Each box spans the interquartile range, horizontal white lines indicate the median and whiskers show the minima and maxima. * indicates a difference at p < 0.05. # indicates a difference at p<0.06.

In the most dorsal population, NCMd, pS6 expression was modulated by stimulus identity (*X^2^* _39)_ = 51.8815, *p* <0.0001), and there was a trend towards an interaction of rearing and stimulus (*X^2^* = 7.6967, *p* = 0.0527; see Fig. 3c). Specifically, while all birds had higher pS6 expression for song playback compared to silence (*X^2^* = 28.2628, *p* <0.0001), females varied in the degree to which pS6 expression was modulated by the bird’s relationship to the singer. In normally-reared females, pS6 expression was greatest when the song belonged to the mate (versus familiar: *p* < 0.0001; versus unfamiliar: *p* = 0.0406; versus silence: *p* < 0.0001). Expression was also high for the song of an unfamiliar male compared to a familiar song or silence (*p* = 0.0109 and *p* < 0.0001, respectively) while the familiar song did not induce significant pS6 expression (*p* = 0.0741 compared to silence). In striking contrast, among song-naïve birds, all song playbacks induced significant expression of pS6 compared to silence (*p* < 0.0001 for all); however, there was no significant variation based on the identity of the song (*p* > 0.5 for all comparisons). A similar pattern of expression was apparent in the NCMp, where there was an interaction between rearing and stimulus identity (*X^2^* = 9.4253, *p* = 0.0241; Fig. 3d). Among normally-reared females, those that heard their mate’s song had significantly higher pS6 expression than those that heard familiar song (*p* = 0.0030), unfamiliar song (*p* = 0.0252), or silence (*p* < 0.0001). Females that heard an unfamiliar song also had greater pS6 expression than those that heard silence (*p* = 0.0107), but there was no difference between those that heard familiar song versus silence (*p* = 0.0609) or unfamiliar versus familiar song (*p* = 0.4632). In contrast, among song-naïve females, there was greater response to song compared to silence for all stimuli (*p* < 0.01 for all). Intriguingly, unfamiliar song elicited the greatest increase in pS6 expression in song-naïve females, which was significantly higher than expression in response to the mate’s song (*p* = 0.0488) and trending towards significantly higher than expression in response to a familiar song (*p* = 0.0550).

In the central region, NCMc, there was a main effect of stimulus but no differences between the rearing conditions (stimulus: *X^2^*_(3, 39)_ = 10.9736, *p* = 0.0119; rearing: *X^2^*_(1, 39)_ = 2.5006, *p* = 0.1138; Fig. 3e). In particular, the response to the mate’s song was significantly greater than either unfamiliar song (*p* = 0.0187) or silence (*p* = 0.0025). Hearing the familiar song also led to increased pS6 expression compared to silence (*p* = 0.0483) while hearing unfamiliar song did not increase pS6 expression over silence (*p* = 0.4473). Finally, in the NCMv, there were no significantt effects of rearing, stimulus identity, or their interaction (*p* > 0.10 for all; Fig. 3f).

### Stimulus identity also modulates activity in the ascending auditory system

In the caudomedial mesopallium (CMM) there were, on average, significantly more pS6-positive cells in normally-reared females compared to song-naïve females (*X^2^* = 5.6584, *p* = 0.0174; Fig. 4f). However, pS6 expression in females from both rearing conditions was similarly modulated by stimulus identity. In particular, pS6 expression was greater in females that heard a song compared to those that heard silence (all pairwise comparisons *p* < 0.0005; Fig. 4g). The increase in pS6 expression was significantly greater in females that heard unfamiliar songs compared to those that heard a familiar song (*p* = 0.0481), but not compared to those that heard the mate’s song (*p* = 0.6863). There was also no significant difference in pS6 expression between females who heard the mate’s song versus familiar song (*p* = 0.1225). These results are similar to previously published results where EGR1 expression was greater, though not significantly, in females that heard unfamiliar songs compared to those that heard the song of a mate (Woolley & Doupe, 2008).

**Fig. 4.**
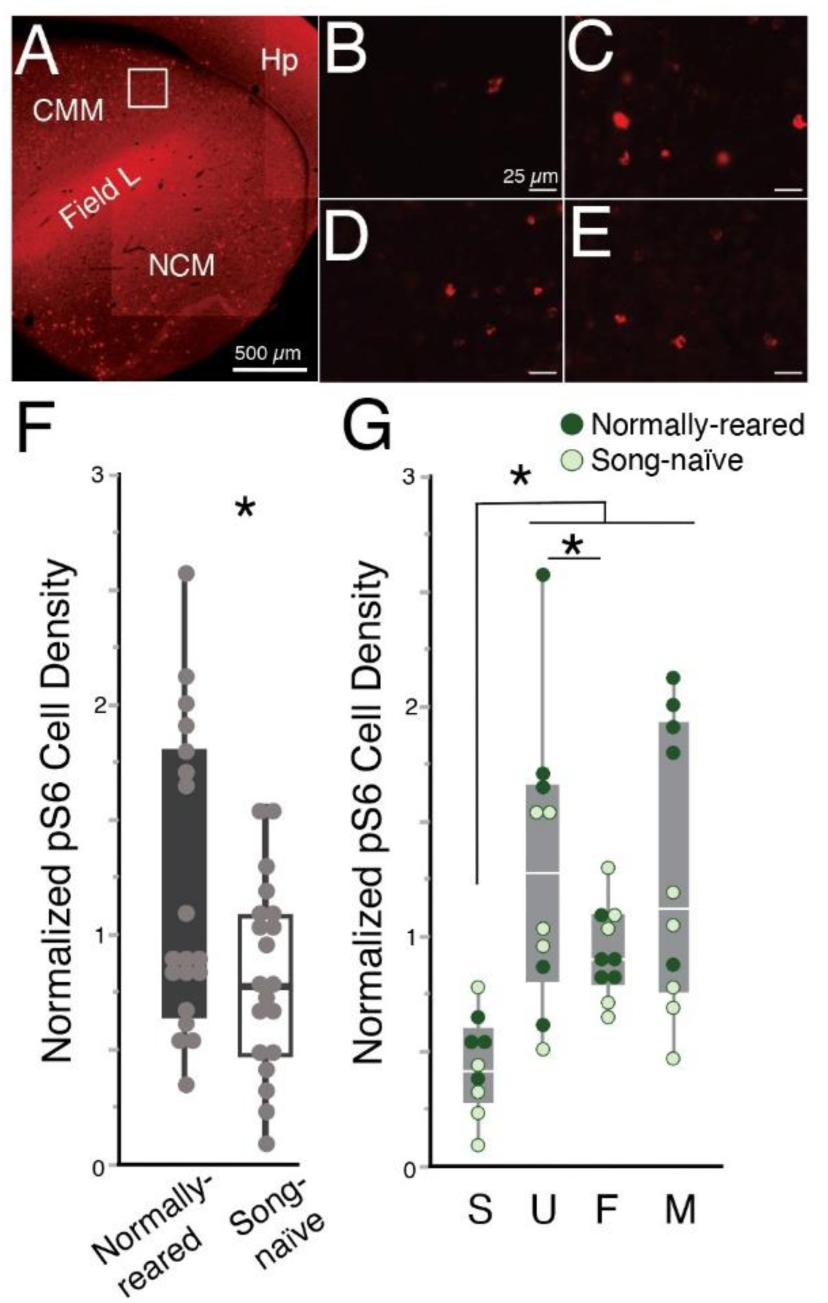
Song-evoked activity in the CMM (A) Widefield view of the caudal auditory forebrain. The white box indicates the location of pS6 counts in the caudomedial mesopallium (CMM). Representative photomicrographs of pS6+ cells (red) in the CMM in (B) silence controls, and birds that heard (C) an unfamiliar song, (D) a familiar neighbor’s song, or (E) their mate’s song. White lines are 25 μm. (F) Box-and-whisker plots showing a higher density of pS6 expressing cells in the CMM of normally-reared females compared to song-naïve females. (G) Box-and-whisker plots showing song evoked activity in females from both rearing conditions. Points are mean counts per individual for normally-reared (closed circles) and song-naïve females (open circles). Each box spans the interquartile range, horizontal white lines indicate the median and whiskers show the minima and maxima.* indicates a difference at p < 0.05.

In the MLd there was no significant effect of rearing (*X^2^*_(1, 39)_ = 1.9019, *p* = 0.1679); however, there was a significant effect of stimulus (*X^2^*_(3, 39)_ = 17.5096, *p* = 0.0006; Fig. 5f). In particular, pS6 was significantly greater in birds that heard unfamiliar song and the mate’s song compared to silence (both *p* < 0.005), while, unexpectedly, pS6 was not significantly increased in females that heard familiar song compared to those exposed to silence (*p* = 0.1832). In females that heard familiar song, pS6 expression was significantly lower than in females that heard the mate’s song (*p* = 0.0090), but not significantly different compared to females that heard an unfamiliar song (*p* = 0.3203).

**Fig. 5.**
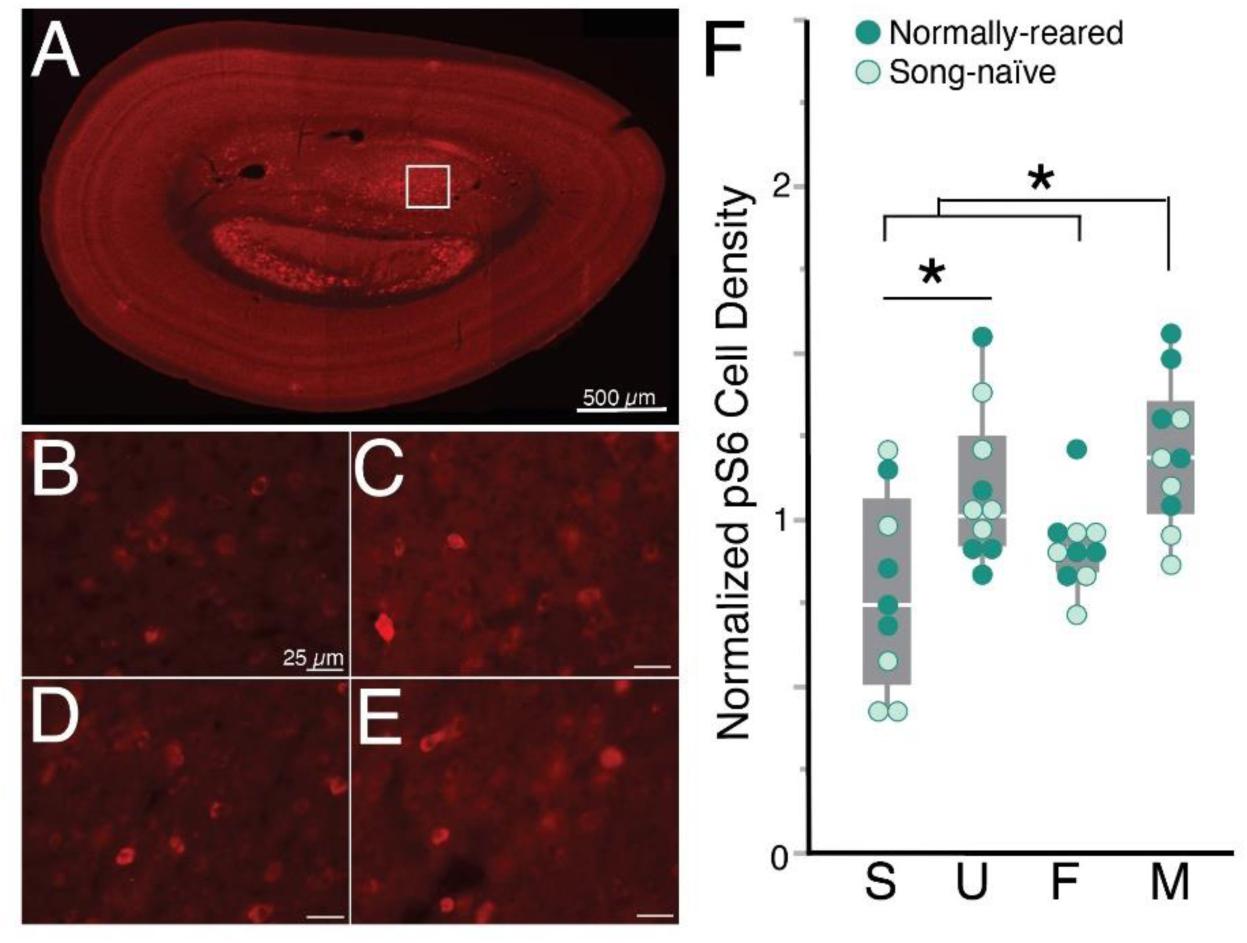
Song-evoked activity in the MLd . (A) Widefield view of the mesencephalic nucleus (MLd). Dashed lines indicate the location of pS6 counts. Representative photomicrographs of pS6+ cells (red) in (B) silence controls, and birds that heard (C) an unfamiliar song, (D) a familiar neighbor’s song, or (E) their mate’s song. White lines are 25 μms. (F) Box-and-whisker plots showing that birds that heard the mate’s song and unfamiliar song had greater pS6 expression than those that heard silence. Points are mean counts per individual for normally-reared (closed circles) and song-naïve females (open circles). Each box spans the interquartile range, horizontal white lines indicate the median and whiskers show the minima and maxima.* indicates a difference at p < 0.05.

## DISCUSSION

Developmental experiences can shape sensory tuning and have sustained, often interactive effects on adult behaviors (Kundakovic & Champagne, 2015; Prounis et al., 2015). While social interactions in adulthood may ameliorate behavioral deficits that arise from aberrant developmental experience, whether these behavioral similarities arise from similarity in the underlying neural circuits is unclear. In zebra finches, a lack of exposure to conspecific song during development has lasting effects on song perception and neural responses and may define or constrain the way these are shaped by adult social experiences. Here, we tested how developmental experience affects adult neural responses to songs that vary in their social valence or familiarity in areas of the avian brain known to be involved in song identification, learning, and preference. While most catecholaminergic regions examined did not show any significant differences, the caudal VTA showed greater activity in the mate song condition, regardless of rearing, an effect that correlates with previous findings of similarities in pair-bonding behaviors and song preferences. In auditory processing areas such as the MLd and CMM, responses also varied by stimulus but not by rearing condition. In contrast, in a region important for song recognition, learning, and memory, the NCM, there were differences in response across subregions. In a central region of the NCM, both normally-reared and song-naïve females had similar patterns of pS6 expression, with the greatest induction of pS6 in response to the mate’s song. In contrast, in two more dorsal subregions there was an interaction of rearing and stimulus: normally-reared females showed greater pS6 expression in response to the mate’s song, while the peak pS6 expression in song-naïve females was in response to the unfamiliar song. Taken together, these data highlight that socially-driven behavioral plasticity in adults may arise through both shared and divergent neural circuits depending on an individual’s developmental experiences.

The VTA is significant in sexual and partner preferences in a diversity of species (Beny-Shefer et al., 2017; Frye & Paris, 2011; Gossman et al., 2024; Nowicki et al., 2020). In voles, the activation of VTA neurons can lead to the formation of pair bonds even in the absence of mating (Curtis & Wang, 2005). Similarly, in zebra finches, peripheral injections of D2 receptor agonists or D2 antagonists can hasten or inhibit the formation of pair-bonds, respectively (Day et al., 2019). More broadly, activity in the caudal VTA is greater for preferred songs (Barr et al., 2021; Barr & Woolley, 2018). In particular, in normally-reared females, cFOS expression in dopaminergic neurons of the VTA is greater for courtship song compared to non-courtship song as well as for normally-tutored versus experimentally tutored song. Early experience modulates courtship song preference, with less consistent preferences for courtship song among song-naïve females. In addition, song-naïve females also do not show an increase in cFOS expression in the cVTA for courtship song compared to non-courtship song. Together, these data hint at a link between song preferences and the level of activity of dopaminergic neurons in the cVTA. Here, we find that, regardless of rearing condition, activation of DA neurons in the cVTA is greater for the mate’s song. Additionally, previous work has also shown that mating ameliorates the preference for courtship song in song-naïve birds (Wall & Woolley, 2024). Whether this means that the activity of the cVTA in response to courtship song in mated, song-naïve females also parallels the activity for normally-reared females is unknown. Future work investigating the degree to which activity in cVTA DA neurons drives or is driven by the formation of the pair bond and the mate’s song preference will lend needed insight into the specific role of the cVTA in song preference learning.

The NCM is a secondary or associative auditory region that has been implicated in song recognition and memory. In adult birds, song from a familiar individual, such as a mate or tutor, or song to which birds were passively exposed through repeated playback induces lower expression of the immediate early gene EGR1 and lower electrophysiological activity within subregions of the NCM compared to a novel song (Chew et al., 1995, 1996; Mello et al., 1995; Mello et al., 1992; Woolley & Doupe, 2008; Woolley & Woolley, 2020). Moreover, NCM lesions and pharmacological manipulations can impair song recognition (Gobes & Bolhuis, 2007; Macedo-Lima & Remage-Healey, 2020; Tomaszycki & Blaine, 2014; Yu et al., 2023). During development, the NCM is implicated in the memorization of the tutor song in juvenile males. Formation of a tutor song memory is associated with changes to neural tuning and the emergence of tutor-song selectivity in the NCM (Phan et al., 2006; Yanagihara & Yazaki-Sugiyama, 2016) and the strength of song learning is correlated with the degree of induction of immediate early genes in response to the tutor’s song (Bolhuis et al., 2000; Bolhuis & Gahr, 2006). Hearing song early in development switches EGR1 expression from constitutively active to inducible (Kudo et al., 2020; Stripling et al., 2001) and disruption of ERK pathways upstream from EGR1 affects tutor song memorization in juvenile birds (London & Clayton, 2008). In female zebra finches, IEG responses in the NCM are also modulated by the attractiveness of song. Across species, females exhibit greater IEG expression to preferred songs compared to less preferred songs (Chen et al., 2017; Gentner et al., 2001; Monbureau et al., 2015). For example, there is greater induction of EGR1 in response to preferred long-bout songs compared to short-bout songs in females starlings (Gentner et al., 2001) and in response to courtship song relative to non-courtship song in female zebra finches (Chen et al., 2017). In zebra finches, silencing or pharmacological manipulation of the NCM alters female song preferences (Barr et al., 2021; Tomaszycki & Blaine, 2014). We hypothesize that the NCM is important in individual recognition and preference and could be a locus encoding the mate’s song memory. Our data support a potential role for the NCM in mate’s song recognition as we find a number of subregions within the NCM, especially in the dorsal and rostral regions, that have higher pS6 expression for the mate’s song over an unfamiliar song or silence.

While our data showing modulation of pS6 expression by singer identity generally support a role for the NCM in mate’s song recognition, we did find region-dependent variation between the rearing conditions in the strength of response to the mate’s song. Developmental song exposure affects both song preferences and the induction of immediate-early genes in response to species-typical song variation in females (Chen et al., 2017). However, song-naïve females learn to prefer the song of a mate in a manner similar to normally-reared females (Wall & Woolley, 2024). Given the ability of both normally-reared and song-naïve females to recognize and prefer the mate’s song, we expected to find regions of the NCM in which responses to the mate’s song was high for both rearing conditions. In the NCMc, pS6 expression was greatest for the mate’s song in females from both rearing conditions. In contrast, pS6 expression in regions dorsal to the NCMc, including the NCMp and NCMd, differed between the rearing conditions. Mate’s song induced the greatest pS6 expression in normally-reared females while in song-naïve females unfamiliar song elicited the greatest pS6 expression. The degree to which this reflects recognition of the mate’s song or a preference for the mate’s song is unclear. However, these data highlight potential topographic differences in the shaping of auditory responses by developmental song exposure as well as in the processing of the mate’s song.

In normally-reared females, across a number of NCM subregions, there was significantly greater pS6 expression for the mate’s song compared to the song of a familiar male. Importantly, our experimental design ensured that, within a cohort of females, one female’s mate’s song was simultaneously the familiar song for a female housed in the same soundbox, and an unfamiliar song for a female in a different soundbox. Thus, the same song was used for all females in a cohort, only the relationship of each individual bird with the singer of the song differed. This design accounts for any male-specific variation in song rate during the two weeks of co-habitation or spectral properties. Females could physically interact with their mate, but could only hear the familiar male. The difference in pS6 expression suggests that the mate’s song is encoded differently than songs that have been merely heard many times. In particular, it seems that the social aspect of the pair-bond, and not a mechanism related to habituation to a familiar stimulus, drives a change in the neural encoding of the mate’s song in normally-reared females. Further investigation of the mechanisms driving memory-formation for a mate’s song versus the habituation to a repeated stimulus will lend needed insight into the mechanisms of auditory recognition and preference.

The NCM is highly dopamine sensitive, with extensive innervation by tyrosine hydroxylase terminals (Van Ruijssevelt et al., 2018; Von Eugen et al., 2020) coming from the ventral tegmental area, including the cVTA and substantia nigra pars compacta (Barr et al., 2021) and high expression of dopamine receptors (Kubikova et al., 2010; Macedo-Lima & Remage-Healey, 2021). Moreover, infusion of dopamine agonists or antagonists during song playback can switch a female’s song preference (Barr et al., 2021). Combined with the results of the current experiment, dopaminergic inputs from the VTA appear to be important in forming song preferences, regardless of how the NCM itself processes song. Further research is needed to confirm whether the VTA neurons that are active when hearing the mate’s song are the same neurons that send projections to NCM, and if so, whether these projections are of equal weight in song-naïve and normally-reared females.

In the MLd there was greater pS6 expression for the mate’s song or an unfamiliar song compared to a familiar song. This result was surprising given previous work that has found that responses in the MLd vary based on the quantity of sound but not necessarily the spectrotemporal features or identity of the song (Dai et al., 2018; Schneider & Woolley, 2010; Woolley & Doupe, 2008). However, unlike the current experiment, in most instances in which the MLd has been studied, birds were presented with different songs. Here, we used the same song but varied the categorial relationship between the female and the singer (i.e. the song could be from a mate or an unfamiliar male). In bats, activity in the inferior colliculus (the mammalian homologue of the MLd) has been found to vary in response to sounds that differ in their categorical representation (Lawlor et al., 2023; Salles et al., 2020, 2024). It is also possible that pS6 highlights a different population of neurons than those that have been previously investigated with either EGR1 or electrophysiology.

The patterns of pS6 induction in both the MLd and the NCM were similar, but not identical to those seen previously with EGR1 expression in response to mate’s song or an unfamiliar song in normally-reared birds (Chen et al., 2017; Woolley & Doupe, 2008). While both pS6 and immediate early genes such as EGR1 lie downstream to similar pathways, there may be both coordination and independence to their activation, as seen with how cFOS, EGR1, and Arc show both overlapping and distinct patters of expression (Leitner et al., 2005; Sockman et al., 2005; Velho et al., 2005; Wada et al., 2006). In NCM specifically, EGR1 expression in is greater in response to a novel song compared to the mate’s song (Woolley & Doupe, 2008), which is contrary to the results we have found in this paper. However, previous work describes pS6 expression in the NCM as clustered and sparse compared to EGR1 expression (Ahmadiantehrani et al., 2018), an effect that our data replicates. Moreover, we saw stimulus specific expression patterns that highlight a diagonal band of activity, parallel to Field L3 and the lamina mesopallialis, especially in females that heard the mate’s song. These results provide a foundation for uncovering the degree to which there is topography or subregion level organization within the NCM.

In the wild, zebra finches live in large, gregarious flocks, and thus are exposed to many individual songs over the course of their lives. Within those flocks, finches form long-term, monogamous male-female pairs, with high partner fidelity, and females form strong, lasting preferences for the song of their mate (Griffith, 2019; Zann, 1996). Our data, finding tuning to the mate’s song in regions of the NCM and VTA fits with the ascendency of pair bonding and mate preference in the species’ life history and ecology (Griffith, 2019). The widespread, preferential response to the mate’s song in the NCM in normally-reared females highlights the significance of this social relationship and raises interesting questions about the mechanisms by which the neural signature for the mate’s song arises over the course of pair-bonding. Moreover, while the similarity in neural activation in the NCMc and VTA between normally-reared and song naïve birds highlights potential subregions that may be important in learning of a preference for a mate’s song, we see lasting evidence of the effects of developmental differences in sensory experience in other NCM subregions, which suggest differences in the neuronal encoding mechanisms of learned song in adulthood that is dependent upon the song exposure or lack thereof received during development. Taken together, these highlight the importance of early life experience in shaping neural functioning throughout adulthood.

## METHODS

### Animals

Zebra finches (N = 67, n = 39 females, n = 28 males, all >90 days post-hatch) had *ad libitum* access to seed, water, and grit and were kept on a 14:10 light:dark cycle. Egg and egg supplements were provided twice a week. Bird care and experimental procedures were approved by the McGill University Animal Care Committee and were performed in accordance with the Canadian Council on Animal Care guidelines.

Birds were reared in one of two conditions. Normally-reared birds were reared with both parents and were exposed to the songs of other males in a colony setting, while song-naïve birds had their father removed 5-7 dph, and were reared by their mother only in sound-attenuating boxes (TRA Acoustics, Cornwell, Ontario). Song-naïve birds remained in the nest until 60 dph, at which point they were moved to a female-only colony setting.

### Experimental Design

Female birds were housed with a male in a cage (10 in. x 10 in.) within a sound-attenuating chamber (“soundbox”). Within the same soundbox, a second male-female pair was housed in a different, adjacent cage. Cages were separated by a visual barrier which allowed birds in each cage to hear but not see or physically interact with birds in the neighboring cage. All pairs were given access to nesting material and an empty dish in which to nest. This setup was repeated across two soundboxes for each rearing condition Thus, for each experimental “batch” there were a total of four soundboxes containing a total of eight pairs of mated birds (four with song-naïve females, four with normally-reared females). We created 5 experimental batches (n = 20 total normally-reared females and n = 19 total song-naïve females). In one exceptional instance, a silent control female was not mated. Birds were housed together for two weeks, which previous research indicates is sufficient time to form a pair bond (Wall & Woolley, 2024).

### Stimulus creation

Prior to mating, males were recorded singing both directed (female-oriented) and undirected song using Sound Analysis Pro (SAP; Tchernichovski et al., 2000) as previously described (Chen et al., 2017; Schubloom & Woolley, 2016; Wall & Woolley, 2024; Woolley & Doupe, 2008). Briefly, directed song was elicited using stimulus females (not used in the study), who were presented to the male for 30 seconds and then removed. Recorded song was then bandpass filtered (300-10 kHz), normalized by the maximum amplitude, and saved as a .wav file (44.1 kHz) using custom Matlab code (Mathworks, Natick, Massachusetts, USA). On average 9 song clips (range 8-10) were selected per male to be used as stimuli. Because within a batch, song-naïve and normally-reared females heard different males, males with similar song durations and numbers of motifs were selected to ensure similar quantities of sound were heard across rearing conditions.

### Stimulus playback

Female birds were separated from their mates overnight. The following morning, lights went on for 30 minutes, beginning one hour and 45 minutes prior to perfusion. Lights were then turned off, and 30 minutes later stimuli were played for the following 30 minutes, also in the dark. Birds were perfused fifteen minutes after playback of the last stimulus.

Birds were organized into 5 immunocytochemical batches, with each batch containing females from both rearing conditions (normally-reared and song-naïve) and all stimulus conditions (mate, familiar, unfamiliar, silence). Specifically, stimuli were such that for each set of four normally-reared and song-naïve females, the song from one male was chosen as the stimulus. This male’s song was played back to three females who each had a different relationship with the male. For one female, the song was from her mate (‘Mate’s song’), for one female the song was from the bird in the adjacent cage within the same sound-attenuating box (‘Familiar song’), and for one bird the song was completely unfamiliar, from a bird in a different sound-attenuating box (‘Unfamiliar song’). The fourth bird in each batch was a silent control, and received silence in place of sound playback.

### Immunocytochemistry and Imaging

After stimulus playback, females were acutely anesthetized via isoflurane inhalation and then transcardially perfused with 12mg heparin in 0.9% saline, followed by 150mL of 4% paraformaldehyde. Brains were removed and stored overnight in 4% paraformaldehyde, after which they were transferred to 30% sucrose for cryoprotection. Brains were sliced in 40 μm sagittal sections and kept in 0.025M phosphate-buffered saline (PBS) with sodium azide before further procedures.

We performed double-label immunocytochemistry to antibodies for phosphorylated S6 (pS6) as a marker of neural activity, and tyrosine hydroxylase (TH) as a marker of catecholaminergic neurons as described previously (Barr & Woolley, 2018; Chen et al., 2017; Fan et al., 2022). Reactions were performed in five immunocytochemical batches with all rearing conditions and stimulus groups represented in each batch, using every third section for each bird. In brief, sections were washed 3×10 minutes with 0.025M PBS, then transferred to blocking for one hour in 5% donkey serum and 0.3% Triton X-100. After blocking, sections were incubated for 48 hours in primary antibody: rabbit-anti-pS6 (1:500 dilution, Cell Signalling Technologies, Danvers, Massachusetts, USA) and sheep-anti-TH (1:1000 dilution, Novus Biologicals, Toronto, Ontario, Canada) at 4° C. After 48 hours, slices were once again washed 2x 30 minutes in 0.025M PBS, and then incubated at room temperature in the secondary antibody: donkey anti-rabbit conjugated to Alexa Fluor 594 (Life Technologies, Eugene, Oregon, USA) and donkey anti-sheep conjugated to Alexa Fluor 488 (Life Technologies). Finally, the sections were washed 3×10 minutes in 0.025 PBS and stored in 0.025 PBS. Sections were mounted on chromium-aluminum subbed glass slides and cover slipped using ProLong Gold Antifade Reagent (Life Sciences, Burlington, Ontario, Canada).

Stained tissue was imaged using Zen microscopy software (Carl Zeiss, Germany) with a 40x objective on a Zeiss Axio Imager upright microscope and an AxioCam MRm Zeiss camera (Carl Zeiss, Germany), using 594 and 488 channels. Exposure times were calculated by auto-determining the exposure of individual areas through Zen, and then averaging the exposure number of each brain region per batch. Exposures > 1000s were removed from the average to avoid overexposing sections that contained large amounts of fluorescence.

Each section was imaged using TH as an anatomical reference (see Fig. 1d), and each brain region was imaged in both left and right hemispheres. We imaged five auditory regions, the caudomedial nidopallium (NCM), caudomedial mesopallium (CMM), caudolateral mesopallium (CLM), the nucleus ovoidalis (Ov), and the nucleus mesencephalicus lateralis pars dorsalis (MLd), and five catecholamine producing regions, the ventral tegmental area (VTA), substantia nigra pars compacta (SN), periaqueductal gray (PAG), and locus coeruleus (LC), as well as the lateral septum (LS) and ventromedial hypothalamus (VMH). The NCM was imaged four times per slice (see Fig. 3b; avg 4.6, range 2-11 sections per region per bird), starting from the most medial slice on which tissue in that region was visible. The most dorsal region was imaged at the very top of NCM (NCMd), the next at the very bottom (NCMv), and the third in the center (NCMc). All of these had one image at 20x zoom per slice, two images per bird. NCMp was imaged as two parts at 40x zoom, the first from where the hippocampus met the NCM. The viewfield was then moved rostrally (towards Field L) until there was no overlap with the previous part and a second part was imaged. Both of these images were counted separately, but analyzed as belonging to NCMp. All subsequent regions were imaged once per slice at 40x zoom. The CMM was imaged where Field L met a ventricle under the hippocampus and formed a triangular shape (avg. 8.6, range 4-11 sections per bird). The CLM was imaged approximately 1.5mm lateral from the midpoint, under the hippocampus and above field L (avg. 7.5, range 5-10 per bird). Ovoidalis was located using autofluorescence to identify the nucleus, approximately 1-1.2mm lateral (avg. 3.9, range 2-4 sections per bird). The MLd was imaged approximately 2.5mm laterally (avg. 7.1, range 4-10 sections per bird). The VTA was imaged twice, in distinct subregions. The first, the rostral VTA, was imaged when the VTA met the edge of the tissue (avg. 4.3, range 2-6 sections per bird). The caudal VTA was measured at the opposite end of the VTA but on the same slices, where it narrowed into the rest of the midbrain (avg. 4.3, range 2-6 sections per bird). The SNc was imaged more laterally than the VTA, when it starts to look rounded on one end and smeared on the other, central in the midbrain (avg. 6.0, range 4-8 sections per bird). The PAG was imaged in roughly the same slices in which the VTA was imaged, along the caudal edge of the midbrain where a ventricle separates the hippocampus (avg. 4.2, range 3-5 sections per bird). The LC was imaged slightly rostral to where the hippocampus joined the midbrain, and identified by a bright cluster of cells (avg. 4.2, range 2-8 sections per bird). The LS was imaged medially, under Field L when it formed a curved or bump shape (avg. 2.8, range 1-4 sections per bird). Finally, the VMH was imaged medially, behind the optic chiasm (avg. 3.1, range 1-4 sections per bird).

Individual cells were counted from raw images using FIJI/ImageJ imaging software (Schindelin et al., 2012; Schneider et al., 2012) by individuals blinded to the conditions of the study. For double-labelled images, cells were counted first on one stain, then separately on the other, before being overlaid to count how many cells were marked on both layers. Counts were then normalized by batch for each area. Each immunocytochemical batch, described in more detail above, consisted of eight birds including females from both rearing conditions (normally-reared and song-naïve) and all stimulus conditions (mate, familiar, unfamiliar, silence). For pS6, the number of cells per image was divided by the size (area) of the image to obtain a count normalized by area. Then, these densities were averaged for each batch, and then the density for each individual image was divided by the batch average. We then took the mean normalized density for each individual. For TH-IR areas, the percentage of active TH cells on each individual image was divided by the batch average of percent active cells. We then took the mean normalized percentage for each individual.

Images used in paper figures were edited for easier visual contrast, and composite images were formed by taking multiple 5x-zoom images and digitally stitching them together to form a complete larger map. However, as mentioned, all data counting used raw images directly with no photomanipulation.

### Statistical analysis

Analysis was conducted using JMP Statistical Processing Software (SAS Institute, Cary, North Carolina, USA). We calculated the density of neurons for each TH-region and batch-normalized the density values as described above to account for variations in immunohistochemical analysis. For TH+ regions, the percentage of TH+ cells colocalizing pS6 was calculated. We performed a Poisson-distributed generalized linear model (GLM) with Log link function. Rearing, stimulus, and their interaction served as independent variables. We used least-squared Tukey’s HSD for all post-hoc tests and set α < 0.05 for all tests unless otherwise noted.

## Notes

### Competing Interest Statement

The authors have declared no competing interest.

### Summary of Updates

Formatting error (location of figures within the text; font of references) and typo corrected.

